# Chronotopic Maps in Human Medial Premotor Cortex

**DOI:** 10.1101/399857

**Authors:** Foteini Protopapa, Masamichi J. Hayashi, Shrikanth Kulashekhar, Wietske van der Zwaag, Giovanni Battistella, Micah M. Murray, Ryota Kanai, Domenica Bueti

**Author notes:** Correspondence to: Prof. Domenica Bueti.

## Abstract

Time is a fundamental dimension of everyday experiences. We can unmistakably sense its passage and adjust our behavior accordingly. Despite its ubiquity, the neuronal mechanisms underlying the capacity to perceive time remains unclear. Here, in two experiments using ultra-high-field 7-Tesla functional magnetic resonance imaging, we show that in the medial premotor cortex of the human brain, neural units tuned to different durations are orderly mapped in contiguous portions of the cortical surface, so as to form chronomaps. The response of each portion in a chronomap is enhanced by preferred and neighboring durations and suppressed by non-preferred durations represented in distant portions of the map. These findings identify duration-sensitive tuning as a neural mechanism underlying the recognition of time and demonstrate for the first time that the representation of an abstract feature such as time can be instantiated by a topographical arrangement of duration-sensitive neural populations.

## Introduction

Time is a particularly elusive dimension of everyday experiences. We cannot see or touch time; nevertheless, we clearly sense its flow and adjust our behavior accordingly. When dancing, our body entrains to the musical tempo. Even without a watch, we can detect when the bus we are waiting for is late.

While a growing body of evidence highlights the contribution of many different brain regions to temporal computations, the neuronal mechanisms underlying our capacity to perceive time remains largely unknown[1][2].

Single-neuron recordings in animals suggest that the encoding of temporal information in the millisecond/second range is achieved via duration tuned mechanisms[3][4][5]. Duration selective cells have been observed in cat’s early visual cortex[5], in cat’s and bat’s primary auditory cortex[6][7], and more recently in the monkey’s medial premotor and prefrontal cortices[3][4][8]. In the human brain, the existence of such mechanisms has been recently suggested by psychophysical studies[9][10] and by a single neuroimaging experiment[11]. Psychophysical studies show that the repeated presentation of a visual stimulus or an auditory rhythm of a given duration (i.e., ‘adaptor’) affects the perceived duration of a subsequent visual stimulus or rhythm (i.e., ‘after-effect’). After-effects are stronger if the temporal distance between the ‘adaptor’ and the judged stimulus is optimal, suggesting the existence of tuning profiles[9][10] where the selectivity is highest for the preferred duration and slowly decays with distance from it. Duration adaptation has also been shown to influence the activity of the inferior parietal lobule (IPL) in the human brain. Neural activity in the IPL is suppressed for stimuli of the same duration and enhanced for stimuli of different durations[11].

However, previous studies, in either the animal or the human brain, have not clarified whether neurons tuned to different durations have an ordered topographical arrangement in duration-sensitive areas of the brain. Whether this ordered arrangement is a specific property of a single or multiple brain regions also remains unknown.

Neuronal tuning and topography are mechanisms widely used in the brain to represent sensory information[12][13], including abstract features like quantities[14]. Showing the existence of a temporal topography could be therefore very important to clarify the computational architecture underlying time perception and to link the representation of time to that of other sensory features like for example stimulus orientation.

## Results

To examine if chronotopic representations exist in the human brain, we used ultra-high-field fMRI at 7T in two distinct experiments. In the first of these experiments (Exp.1) we measured brain activity while participants (N=11) decided whether the second stimulus (S2) of a pair was longer or shorter than the first one (S1, see Figure 1A). In this experiment, we used 4 different duration ranges (i.e., S1 equal to 0.2, 0.4, 0.6 and 1 s). Stimuli were visual gratings (i.e., Gabor patches) varying in both orientation and duration. Orientation changes were task irrelevant (see Materials and Methods for details).

**Figure 1.**
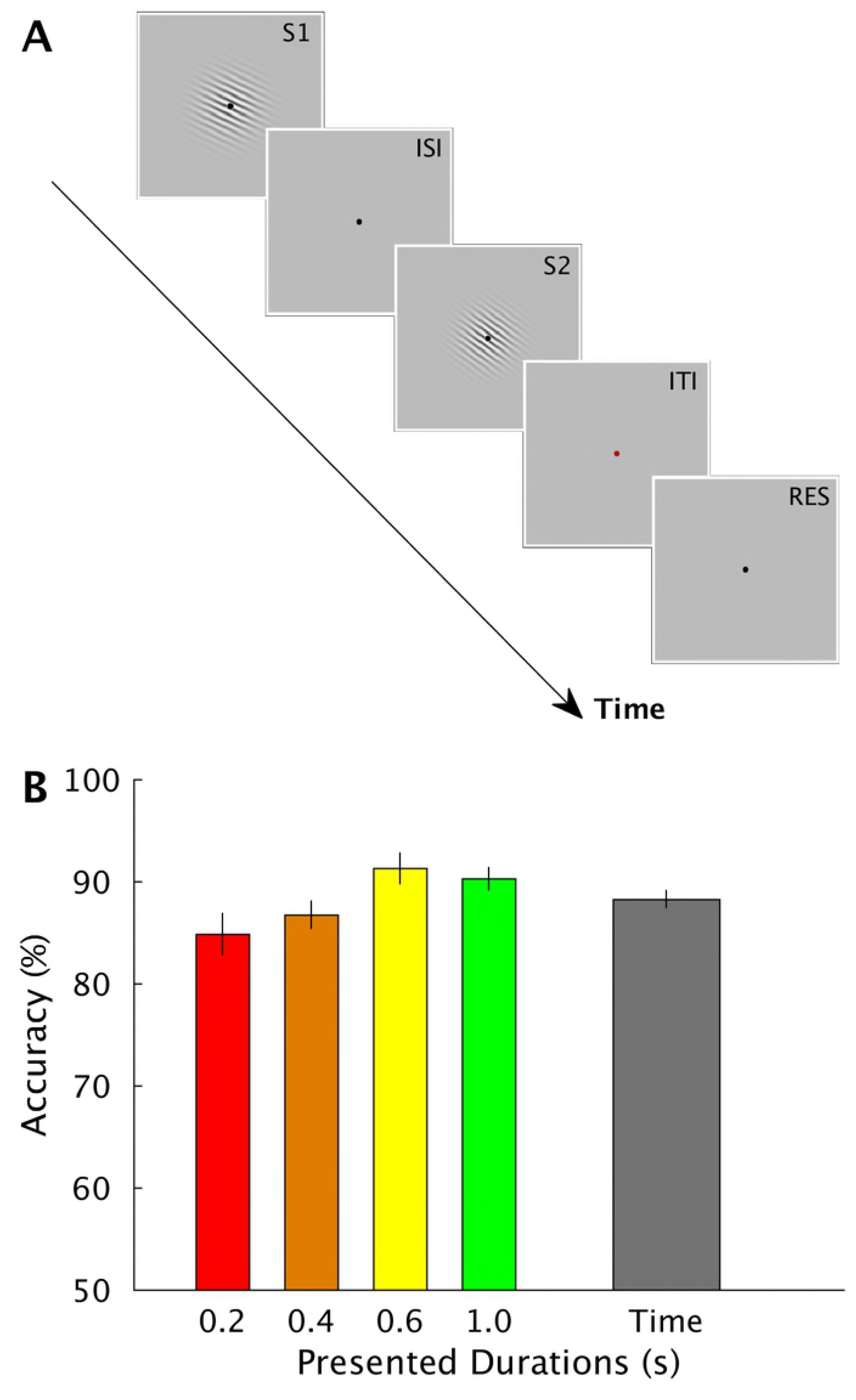
Stimulus sequence and behavioral results of Exp.1. (A). Schematic representation of the stimulus sequence in a trial of Exp.1. In each trial a standard (S1) and a comparison duration (S2) were presented in sequence. S1 could be one of four different durations (0.2, 0.4, 0.6, and 1 s). S2 could be either shorter or longer than S1 (Weber ratio was set to 0.4). Stimuli were sinusoidal Gabor patches varying in orientation. Orientation changes were task irrelevant. Participants were asked, by pressing one of two response keys, to judge whether the duration of S2 was shorter or longer than S1. (B) Group average (N=11) of percentage of accuracy in the time task plotted separately for each of the four durations and as a mean of them (‘overall accuracy’, rightmost bar). Error bars are standard errors.

<u>Please Figure 1 here</u>

Behavioral data indicate that participants performed equally well in all tested durations (see Figure 1B). Proportion of correct responses for each S1 duration condition (i.e., 0.2, 0.4, 0.6, and 1 s) were 85.1 ± 7.1 (mean ± standard deviation), 87.0 ± 4.9, 91.5 ± 5.4 and 90.6 ± 4.1 %, respectively. Overall accuracy was 88.6 ± 3.7 %. Although a one-way repeated measures ANOVA with within-subject factor of S1 durations showed a significant main effect (F_3,30_ = 4.824, p < 0.05), pair-wise post-hoc tests showed no significant difference between the different combinations of S1 durations (all p’s > 0.05, Bonferroni-corrected for multiple comparisons).

For the analysis of Exp.1, we used separate regressors for each of the 4 different duration ranges. The regressors of our General Linear Model (GLM) modeled the offsets of the first intervals and were convolved with the canonical hemodynamic response function (HRF). We used event offset because it was the moment when the duration of a stimulus became available to participants. We first identified the regions associated with the presentation of the four S1 durations together. As expected from previous neuroimaging findings[15][16], these regions were visual, parietal and frontal cortices (see Supplementary Figure 1 and Supplementary table 1). We then focused on the identification of the brain regions that were maximally activated for each specific S1 duration and that clearly showed a topographical arrangement of duration selective voxels.

Figure 2 upper panel shows the group-level significant clusters computed for each of the 4 duration ranges in the temporal task (p_FWE_-cluster level < 0.05, corrected for multiple comparisons across the whole brain). Each color codes the cluster of voxels that was classified, according to a winner-take-all procedure, based on t-statistic maps, as maximally responsive to each of the different duration ranges. The color scale ranges from red, corresponding to voxels responsive to the shortest duration (0.2 s), to green, the voxels maximally responsive to the longest duration (1 s).

**Figure 2.**
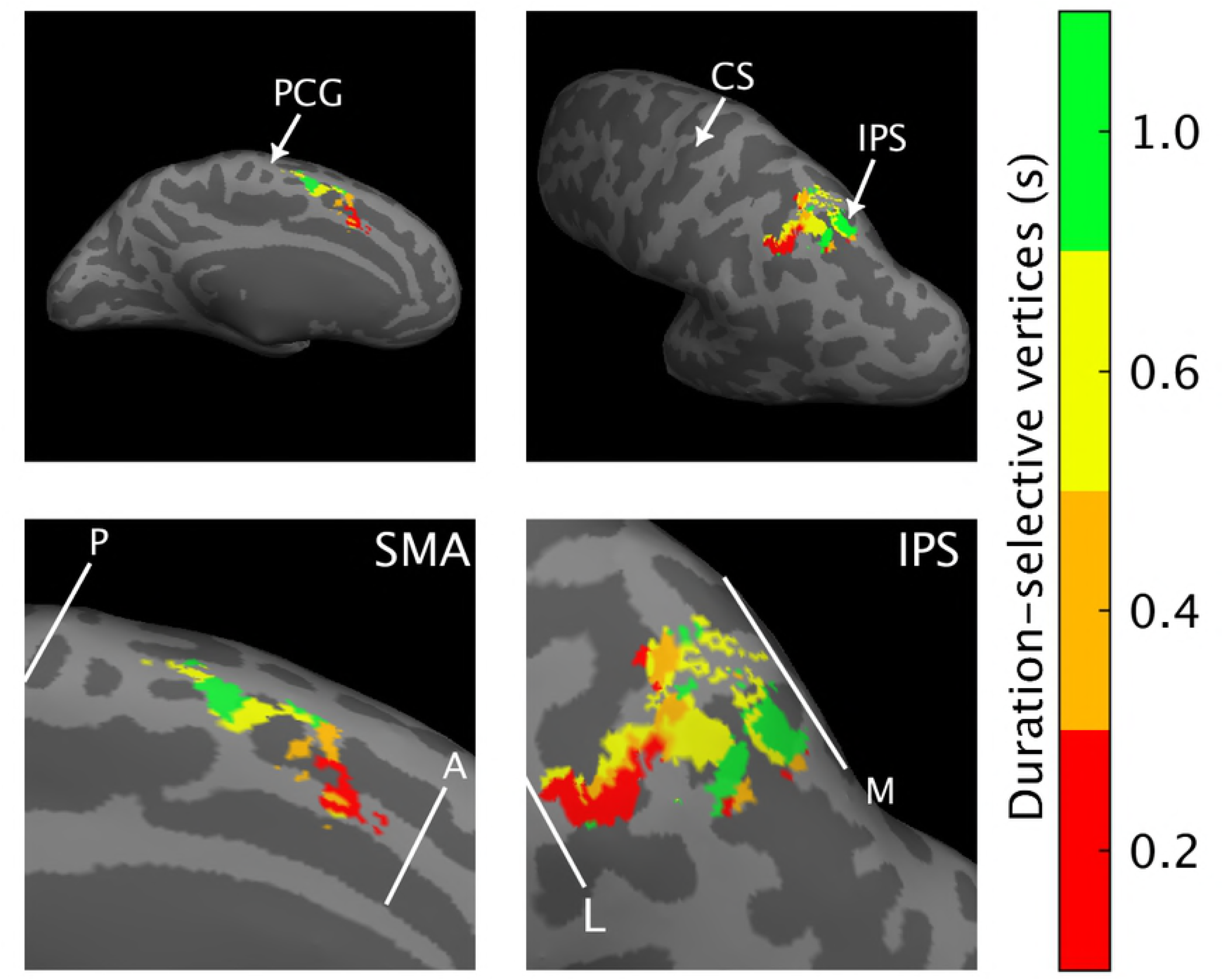
Group-level fMRI results of Exp.1. Medial and lateral view of the left (L) hemisphere with the group-level statistical results (N=11) overlaid on the inflated Dartel-11 template. The figure shows the cluster of vertices (i.e., voxels projected onto the brain surface) classified, according to a winner-take-all procedure based on statistical t-maps, as maximally responsive to each of the four S1 durations (0.2, 0.4, 0.6, and 1 s). Each color codes a different label; the color scale goes from red (shortest S1) to green (longest S1). Statistical threshold for t-maps was set to p_FWE_< 0.05 cluster level corrected for multiple comparisons across the whole brain. Duration selective vertices were found in SMA (leftward panel) but also in the Intraparietal Sulcus (IPS) The durations of the colorbar are red= 0.2, orange=0.4, yellow=0.6, and green= 1 s. Legend: PCG= precentral gyrus, CS= central sulcus, A=anterior, P=posterior, L=lateral, M=medial.

<u>Please Figure 2 here</u>

As indicated by the gradual changes of color in Figure 2, we found a topographic organization of duration sensitive voxels in the supplementary motor area (SMA, see leftward panels) and in part of the intra-parietal sulcus (IPS) of the left hemisphere (see rightward panels). In SMA, this progression was in the rostro-caudal direction with voxels sensitive to the shortest duration located in the anterior premotor cortex and those sensitive to the longest duration in the posterior part. In the IPS the progression was in the lateral-medial direction i.e., voxels maximally responsive to the shorter duration were closer to the lateral border of the map compared to those sensitive to the longest duration.

To quantitatively assess the spatial distribution of duration-selective voxels in SMA and IPS during the temporal discrimination task we analyzed both volumetric and surface data of each individual subject (see Materials and Methods for details) and we chose to look at the spatial progression of the chronomaps by using multiple indexes.

At the surface level, for each subject and each duration selective cluster of vertices (i.e., voxels projected onto the brain surfaces) we calculated the weighted relative distance (wRD) from the posterior and the lateral border of the chronomap for respectively SMA and IPS (see Materials and Methods for details). Borders of the maps were identified in each individual subject.

Figure 3A shows for the left SMA, the median, the quartile range and the fitted slope of wRD of the group (for individual data see Supplementary Figure 2).

**Figure 3.**
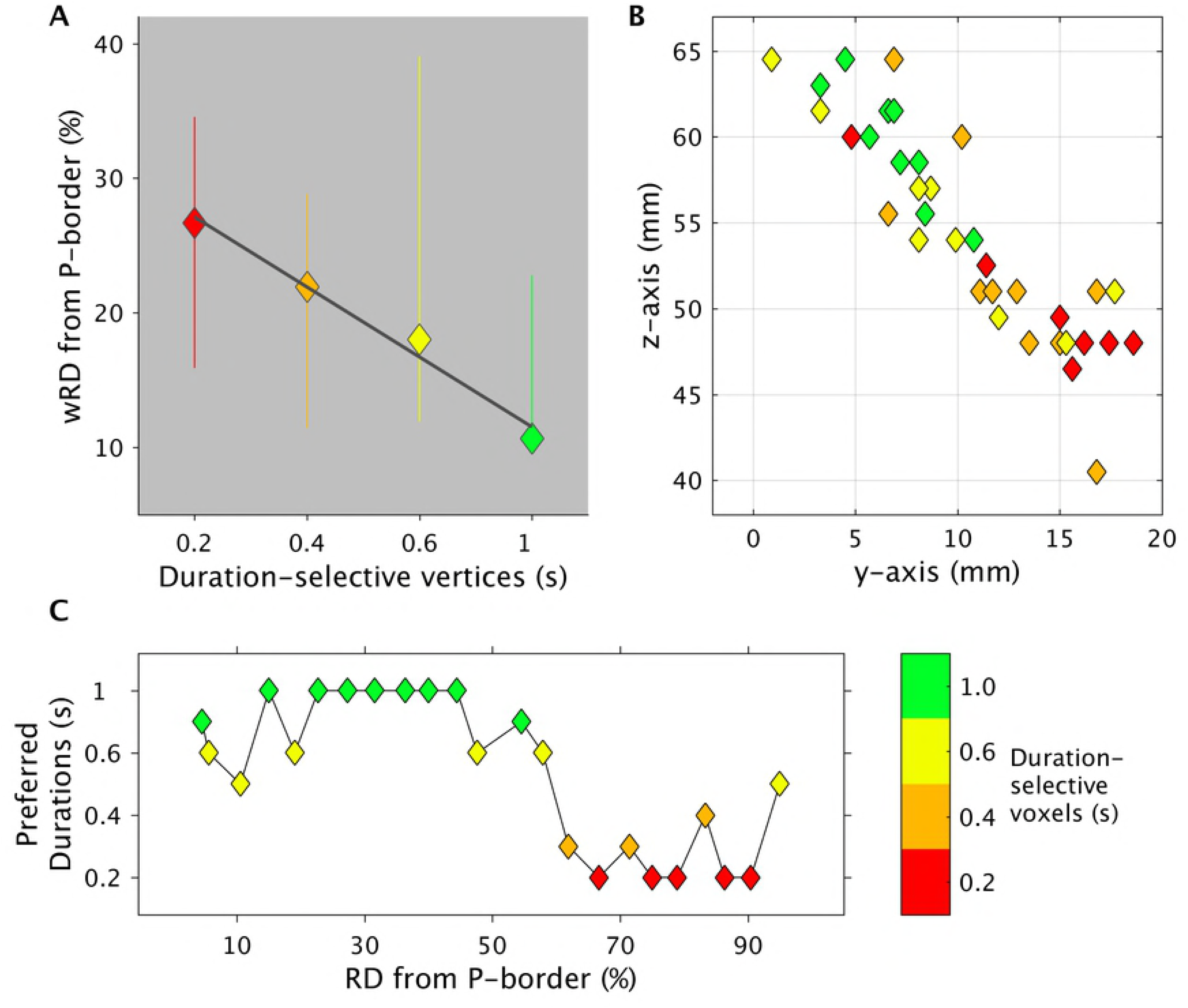
Spatial progression of SMA chronomaps in Exp.1. Panel (A) shows for each duration selective vertex the group median (colored diamonds), the quartile range (vertical bars) and the fitted slope of the “weighted Relative Distance (wRD)” from the posterior border (P) of the chronomaps. wRD were first computed for each individual subject on chronomaps overlaid on flattened surfaces in participant’s native space. The posterior border was chosen to be close the precentral gyrus. (B) weighted centroids (wCntrs) for duration selective voxels in SMA. 2-D projection of wCntrs in the x-y plane. wCntrs are color-coded according to duration selectivity. The color scale goes from red (shortest S1=0.2 s) to green (longest S1=1 s). Different colors indicate voxels with different duration selectivity; diamonds with the same color represent the different subjects (n=11). This last number could change because not all subjects have the full range of duration selective voxels. (C) Group average of preferred duration (y-axis) of voxels lying at different distances (x axis RD = relative distance) from the posterior border of the chronomaps. Legend: P=posterior, wRD =weighted relative distance.

<u>Please Figure 3 here</u>

The plot shows, as expected from the visual inspection of the group-level brain map, that the distance from the posterior border of the SMA is longer for vertices responsive to the shortest duration (0.2 s) and becomes progressively shorter for vertices responsive to the longer duration range. This progression was also present in the majority of the subjects (for individual maps see Supplementary Figure 2) as revealed by the statistically significant analysis of the wRD slopes (Wilcoxon test p=0.017).

To confirm the spatial progression of SMA chronomap, we also identified, for each individual volumetric map, the duration preferred by the majority of the activated voxels that laid at different distances from the posterior border of the chronomap (individual chronomaps were parceled in volumetric bins of 1.5 mm width, for details see Materials and Methods). The relative distance from the posterior border of these preferred durations for the group is shown in Figure 3C. As seen previously, the shorter the distance from the posterior border, the greater the number of voxels preferring the longer duration ranges (diamonds in colder colors). The greater the distance from the posterior border, the greater the number of voxels preferring the short duration ranges (diamonds in warmer colors). A very similar result is shown in Figure 3B where we plot for each subject the weighted centroids of each duration selective cluster. Within the SMA, the centroids of the shortest duration selective cluster (red diamonds) are generally located anteriorly compared to the centroids of the longest duration selective cluster (green diamonds).

In the IPS, the topographical arrangement of voxels (i.e., from lateral to medial for short to long durations), was apparent at the group level, but it was less consistently observed at the single-subject level (see Supplementary Figures 3). Indeed only 5 out of 11 subjects showed the appropriate spatial distribution of duration selective voxels. Moreover, when we looked at the wRD, there was no statistically significant effect of the slope (Wilcoxon test p=0.737, see Supplementary Figure 4).

**Figure 4.**
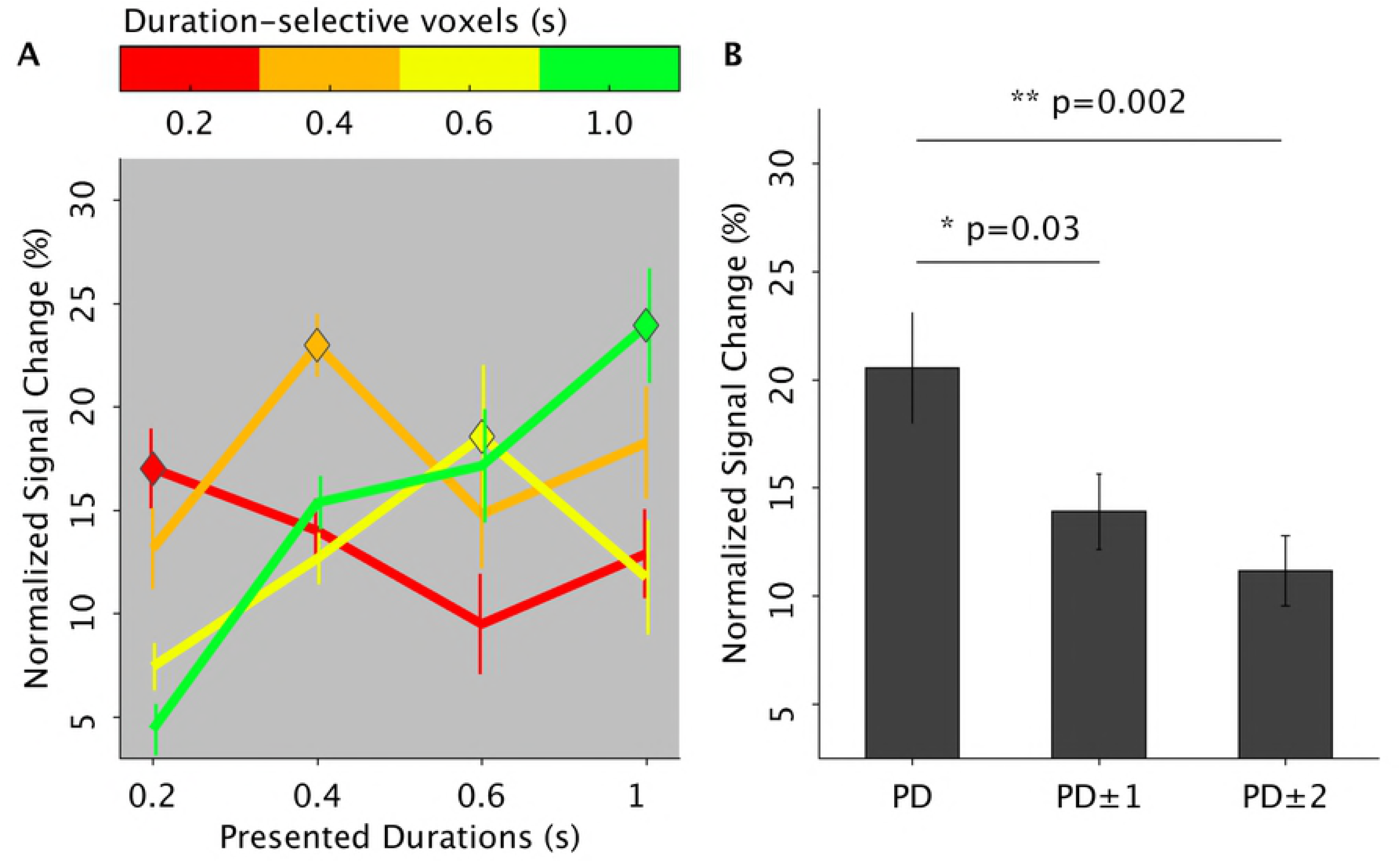
Duration tuning of Exp.1. (A) Group average of normalized BOLD responses (y axis) of duration selective voxels (different lines are different duration selective voxels) for preferred and non-preferred durations. In the x-axis are the 4 presented durations. The BOLD signal in duration selective voxels is aligned to the presentation timings of the different duration ranges (i.e., 2^nd^ volume after S1 offset). The colored diamonds represent the point in time where the hemodynamic response of duration selective voxels matched the presentation timing of the appropriate duration (e.g., red-labeled voxels when the shortest S1 duration is presented). The color code is as in Figure 2. (B) Normalized BOLD response to preferred (PD), neighboring (PD±1) and distant durations (PD±2) averaged across subjects and duration selective voxels. Error bars are standard errors.

To examine the response tuning of the voxels sensitive to a given duration range, we next looked at the change of the hemodynamic response of these voxels for preferred and non-preferred durations. Figure 4 shows the hemodynamic response of duration sensitive voxels for the left SMA. As shown in panel A, for all duration selective clusters (i.e., colored lines), we observed a modulation of the presented durations on the BOLD response. Specifically, the hemodynamic response peaked during the presentation of the preferred duration (PD, see the diamonds in the plot) and slowly decayed for durations distant from the preferred one (PD vs PD±1 p<0.03; PD vs PD±2 p<0.002).

Similar results were obtained in the IPS (Supplementary Figure 5) where the BOLD response was enhanced for preferred (PD) and neighboring (PD±1) durations (PD vs PD±1, p<0.009) and suppressed for durations far (PD±2) from the preferred one (PD vs PD±2 p<0.005).

**Figure 5.**
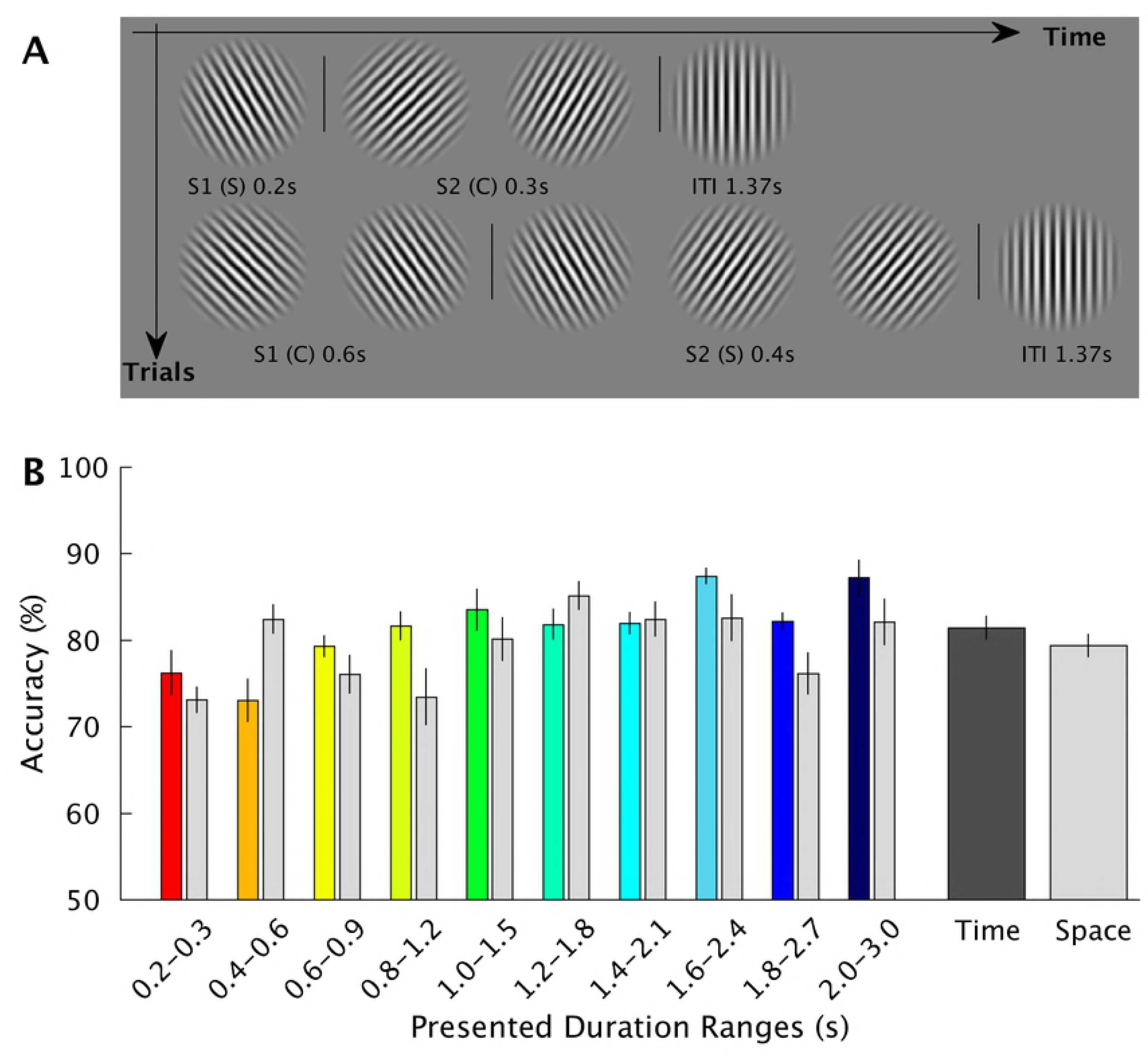
Stimulus sequence and behavioral results of Exp.2. (A) Schematic representation of the stimulus sequence in a trial (Exp. 2). Within a trial, the sequence of orientation changes was fixed and was always leftward first, rightward second and vertical last. Within the two ‘main orientations’ (left and right) the grating continuously changed its orientation at a rate of 5 Hz and the range of changes was between 30° and 45°. Participants were asked to discriminate which of the two ‘main orientations’ (leftward or rightward) was displayed for longer time (time task) or to judge which one of them underwent the biggest change (space task). S is standard and C the comparison duration (there were 10 standard durations ranging from 0.2 to 2 s, in step of 0.2 s). The presentation order of S and C was randomized and counterbalanced across trials (in half of the trials S1 was a standard, in the other half it was a comparison duration). The comparison duration was 50% of the standard. The vertical orientation signaled the time to make the response (by pressing one of two response keys on a keypad) and it was also the inter-trial-interval (1.37 s). (B) Average percentage of accuracy (N=10) in the time and space task plotted separately for each of the 10 pairs of durations and as a mean of them (rightmost plot for time and space tasks). Error bars represent standard errors.

<u>Please Figure 4 here</u>

In order to assess the robustness of Exp.1’s results, we ran an additional experiment (Exp.2, N=10) in which we used a similar temporal discrimination task of visual stimuli (i.e., participants judged which of the two successive visual stimuli (S1 and S2) lasted for longer time). Visual stimuli were Gabor patches changing in orientation (see Figure 5A). In Exp.2 we introduced 3 main changes compared to Exp.1.

First, we used a broader range of durations, spanning from 0.2 to 3 s. Second, we used a method of stimulus presentation that was highly regular, i.e., different durations ranges were presented sequentially. We used pairs of stimuli (S1 and S2) varying in duration. In different pairs we tested different duration ranges e.g. S1=0.2 versus S2=0.3s in one pair and S1=0.4 versus S2=0.6 s in a different pair (see Figure 5A). In each pair we had a standard (T) and a comparison duration (T+ΔT); in half of the trials the standard duration was S1 in the other half it was S2. The pairs were presented in a sequential manner as to form cycles (i.e. a cycle is a series of trials (N=10) where we tested 10 duration ranges). In ascending cycles, we progressed from the shortest to the longest pair of stimuli, in descending cycles it was the opposite.

This design allowed us to evaluate whether there was a gradual spatial shift in cortical activation as the stimulus duration changed.

Third, in addition to the temporal discrimination task, participants performed a non-temporal task in which they judged the spatial orientation of the same visual gratings.

This task was included to evaluate the task-dependency of chronotopic representations.

In Exp.2, S1 and S2 stimuli were defined by different orientations (see Figure 5A and Materials and Methods for full details of the tasks). S1 was leftward and S2 was rightward oriented. While keeping their main orientation, both S1 and S2 slightly changed their angular orientation. In the temporal task participants judged which stimulus orientation was maintained for longer time, whereas in the spatial task they judged which orientation underwent the biggest angular change.

Behavioral data inside the MRI scanner did not reveal any significant performance differences across the different durations (see Figure 5B, main effect of duration F_9_ = 1.303, p = 0.289) and the two tasks (main effect of task F_1_= 0.309, p = 0.592, interaction effect: F_1,9_ = 0.539, p = 0.842).

<u>Please Figure 5 here</u>

At the brain level, based on Exp.1 results, we focused on the identification of chronomaps in both SMA and IPS (for the details on the two Regions of Interest -ROIs see Material and Methods). Given the cyclical presentation of events in the experimental design, data were analyzed with the population Receptive Field method (pRF). pRF is an fMRI method of data analysis that is used to map response selectivity to any type of stimulus feature (e.g. the spatial position of a visual object [17][18]). The idea behind pRF is that neuronal receptive fields are a form of tuning functions. As pRF models we used a one-dimensional Gaussian curve with 2 parameters: *µ*, the stimulus duration and σ, the spread of the pRF. For the pRF modelling we used the offset of all S1 durations, no matter whether S1 was a standard or a comparison duration. This procedure led to the identification of 17 durations (ranging from 0.2 to 3 seconds). For each time point of the fMRI timeseries the overlap between the Gaussian tuning models and the presented stimulus profiles were estimated (see Material and Methods for more details).

Figure 6 shows for the group the projection on the cortical surface (medial part of BA6) of the estimated *µ* parameter. Different colours represent vertices (i.e., voxels projected onto the cortical surface) selective to different duration ranges (i.e., vertices with different estimated *µ).*

**Figure 6.**
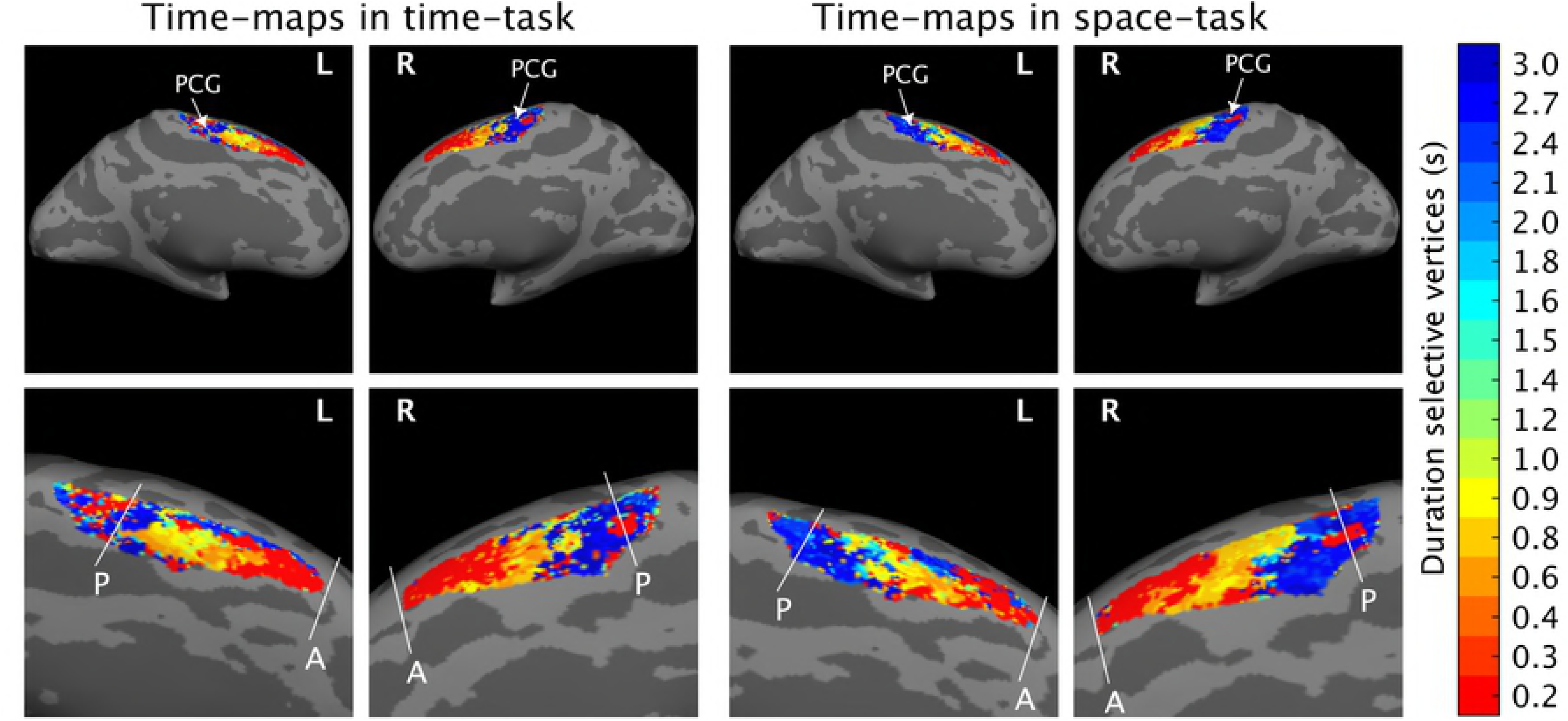
pRF Group-level results of Exp.2. Here we show the projection on the cortical surface (medial part of BA 6) of the estimated *µ* parameter. Different colours represent vertices (i.e., voxels projected onto the cortical surface) selective to different duration ranges (i.e., vertices with different estimated *µ*). We show the results of the group (average of 10 subjects) for the 17 estimated *µ*. The 17 *µ* are the 17 durations presented in the 10 different trial types (S1 duration each time it was either a standard or a comparison duration). The color scale goes from red i.e., shortest duration (0.2 s) to dark blue i.e., longest duration (3 s). The white lines give an example of the map borders as they were drawn to estimate the weighted Relative Distance in the individual subjects. On the left-hand side, time maps in time task, on the right-hand side time maps in the space task. Legend: L=left, R=right, PCG= post central gyrus, SMA= Supplementary Motor Area, A=anterior, P=posterior.

<u>Please Figure 6 here</u>

As indicated by the gradual changes of color in brain activations shown in Figure 6, we found a topographic organization of duration sensitive voxels in the left SMA replicating the results of Exp. 1. In addition to the first experiment, here we observed chronotopic maps for a broader range of durations, in both the left and the right hemisphere and for both temporal and spatial task (see leftward and rightward panels of Figure 6).

As in Exp. 1, this progression was in the rostro-caudal direction within the SMA, with voxels sensitive to the shorter duration (voxels in warmer colors) located in the anterior and those sensitive to the longer duration (voxels in colder colors) in the posterior SMA.

In analogy with Exp.1 we looked at the spatial progression of chronomaps using 3 distinct indexes: wRD, preferred durations and weighted-centroids (see Materials and Methods for details).

Although at a visual inspection (see Figure 6) of the group level results, chronotopic maps seemed to be present in both hemispheres and for both tasks, the analysis on the wRD revealed that in both tasks, only vertices of the right hemisphere showed a very clear spatial progression. Indeed, only in the right hemisphere of both tasks, voxels selective to the longest duration were significantly closer to the posterior border compared to vertices sensitive to the shorter durations (see Figures 7-8 panel A for the temporal and the spatial task respectively; Wilcoxon test on the wRD slope: time task, right hemisphere p<0.001, left hemisphere p=1, space task, right hemisphere p<0.001, left hemisphere p=1). Indeed, only in the right hemisphere this spatial progression was consistently present for the majority of the tested subjects (for individual maps see Supplementary Figures 6-9. For left and right SMA in the temporal task see Supplementary Figures 6 and 7. For the left and right SMA in the spatial task see Supplementary Figures 8 and 9).

**Figure 7.**
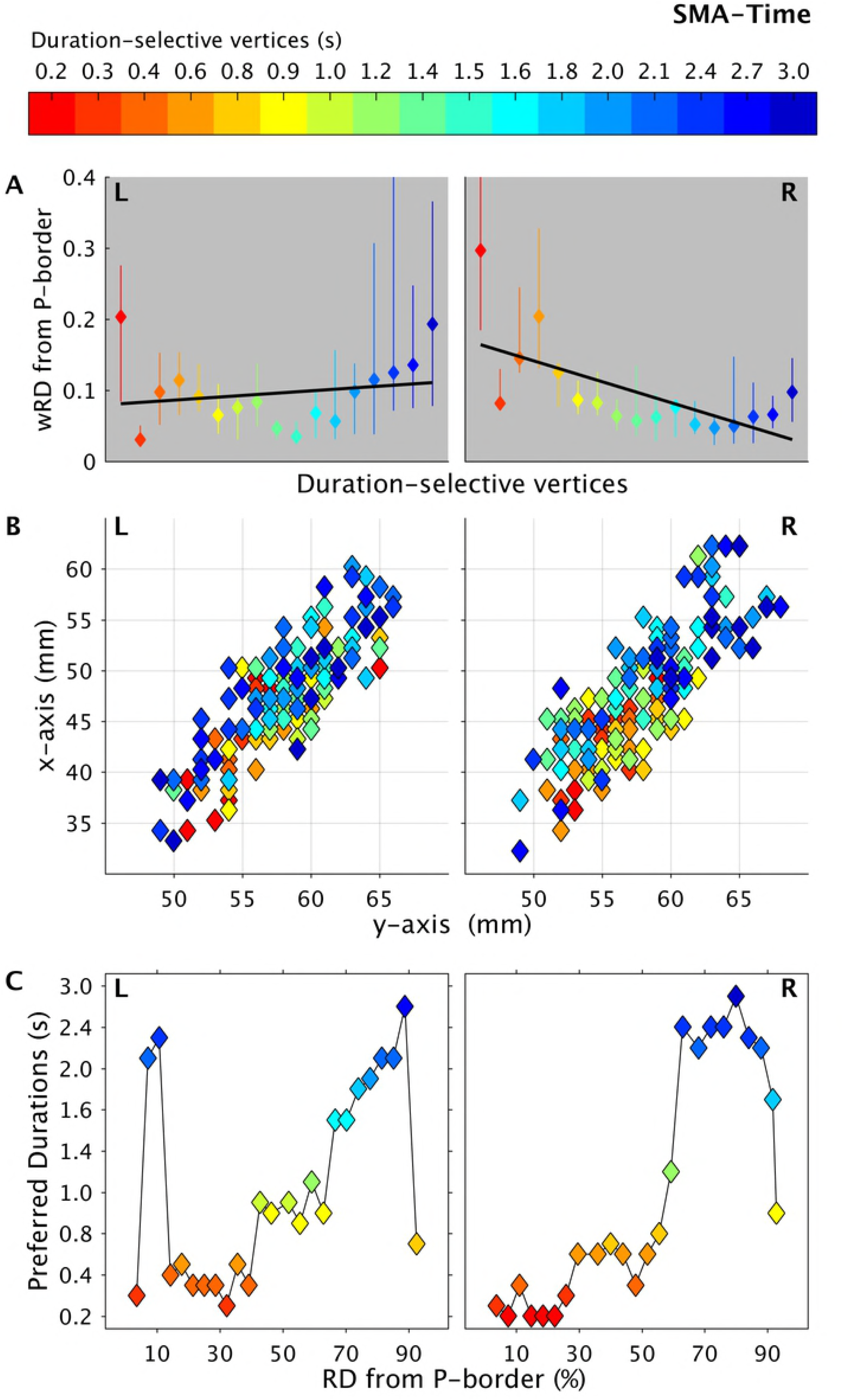
Spatial progression of left (L) and right (R) SMA chronomaps in Exp.2 during the time task. (A) show for each duration selective vertex the group median (diamonds), the quartile range (vertical bars) and the fitted slope of the “weighted Relative Distance (wRD)” from the posterior border (P) of the chronomaps. wRD were first computed for each individual subject on chronomaps overlaid on flattened surfaces in participant’s native space. The posterior border was chosen to be close the precentral gyrus. (B) 2D projection of weighted centroids (wCntrs) in the x-y plane for duration selective voxels in SMA. wCntrs are color-coded according to duration selectivity. The color scale goes from red (shortest duration 0.2 s) to dark blue (longest duration 3s). Different colors indicate voxels with different duration selectivity; diamonds with the same color are the different subjects (n=10). This last number could change because not all subjects have the full range of duration selective voxels. (C) Group average of preferred duration (y-axis) of voxels lying at different distances (x axis RD = relative distance) from the posterior border of the chronomaps. Legend: P=posterior, wRD =weighted relative distance.

**Figure 8.**
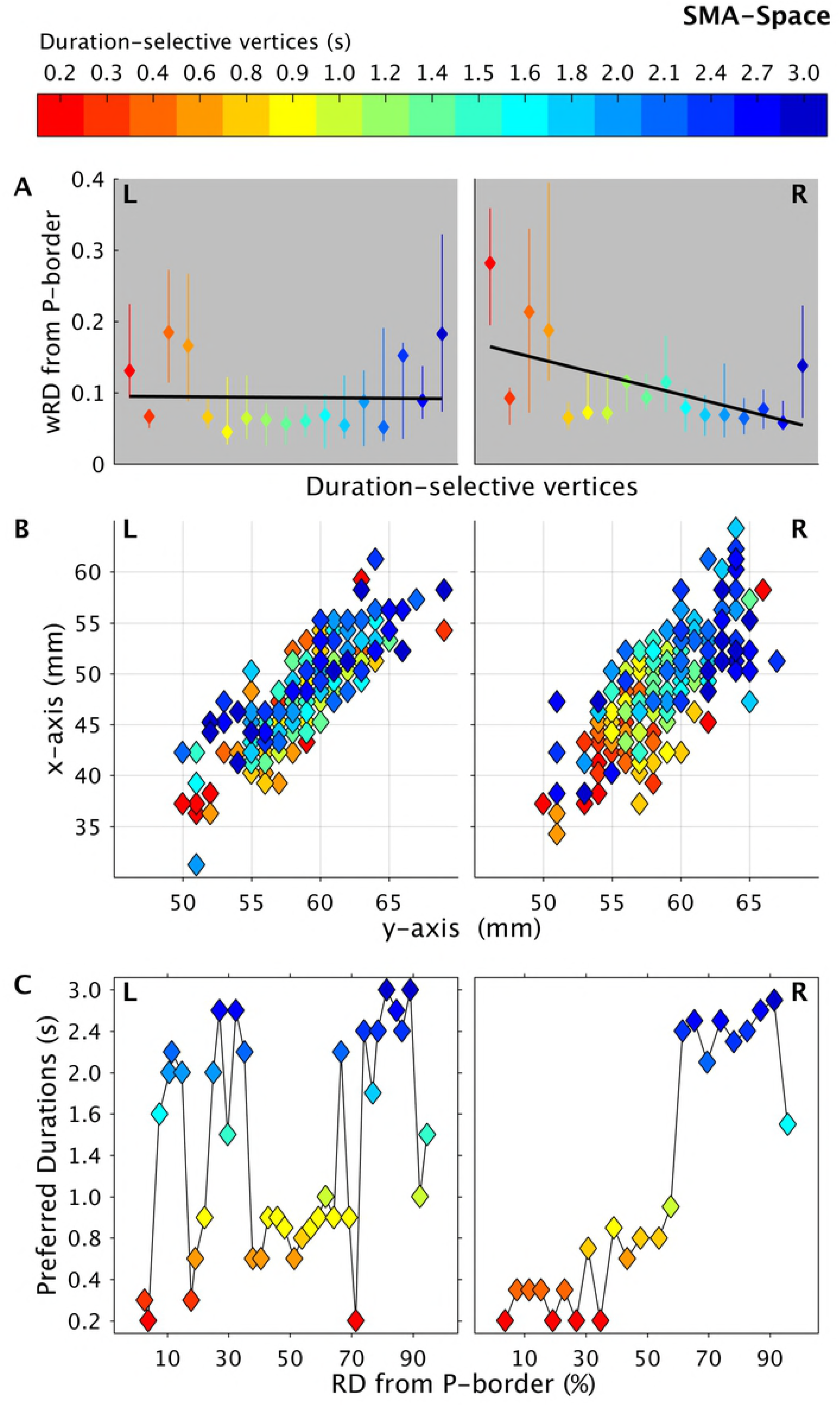
Spatial progression of left (L) and right (R) SMA chronomaps in Exp.2 during the space task. (A) show for each duration selective vertex the group median (diamonds), the quartile range (vertical bars) and the fitted slope of the “weighted Relative Distance (wRD)” from the posterior border (P) of the chronomaps. wRD were first computed for each individual subject on chronomaps overlaid on flattened surfaces in participant’s native space. The posterior border was chosen to be close the precentral gyrus. (B) weighted centroids (wCntrs) for duration selective voxels in SMA. 2-D projection of wCntrs in the x-y plane. wCntrs are color-coded according to duration selectivity. The color scale goes from red (shortest duration 0.2 s) to dark blue (longest duration 3s). Different colors indicate voxels with different duration selectivity; diamonds with the same color are the different subjects (n=10). This last number could change because not all subjects have the full range of duration selective voxels. (C) Group average of preferred duration (y-axis) of voxels lying at different distances (x axis RD = relative distance) from the posterior border of the chronomaps. Legend: P=posterior, wRD =weighted relative distance.

**Figure 9.**
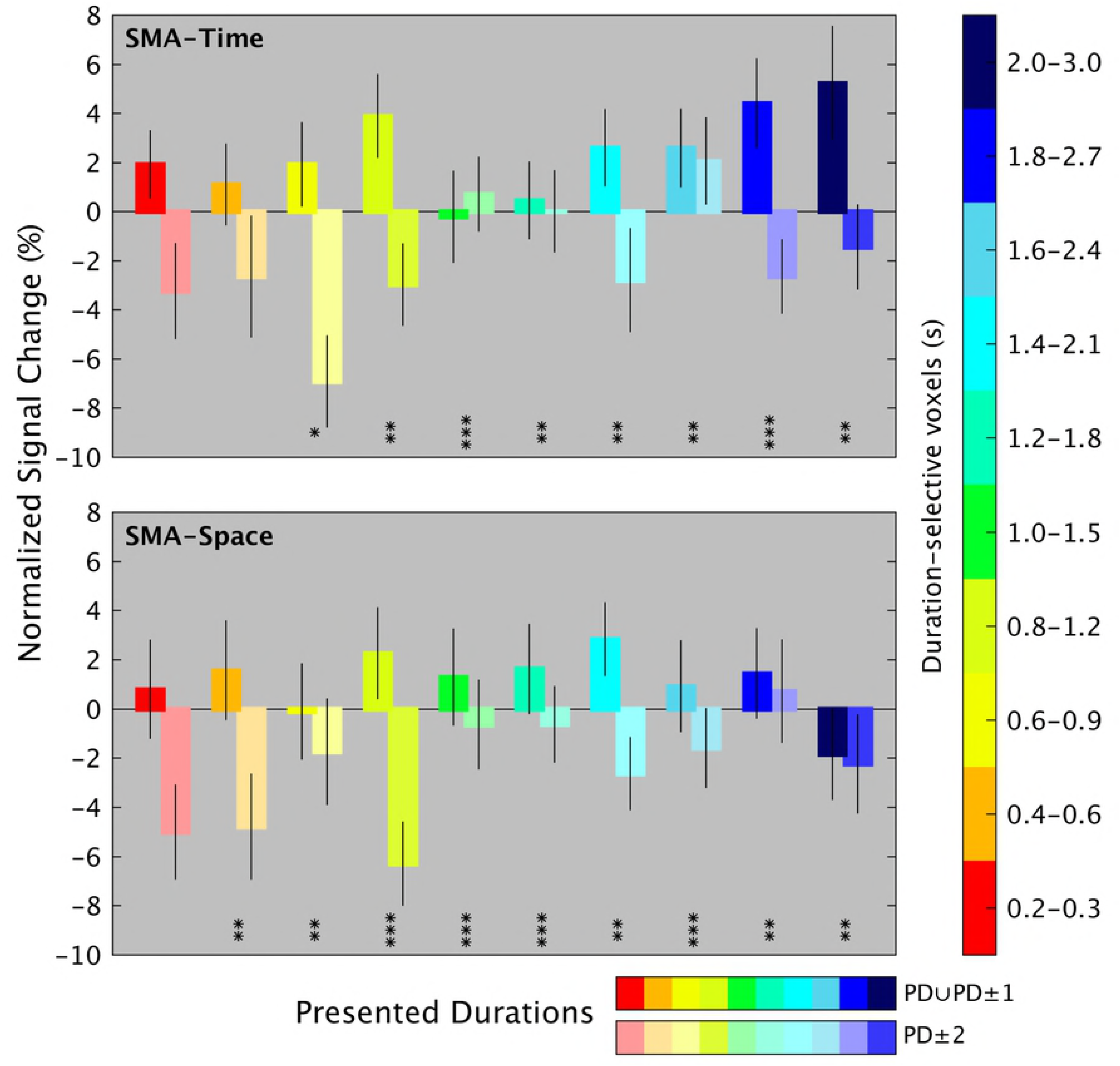
Duration tuning of Exp.2. Group average of normalized BOLD responses of duration selective voxels (colored bars, y axis) for preferred (PD) and neighboring (PD ±1), durations (PD ∪PD±1, bars with darker shades), as opposed to distant non-preferred durations (PD ±2, bars with lighter shades). Asterisks indicate statistically significant difference at a Wilcoxon rank sum test between PD ∪ PD±1 and PD ±2 at *p<0.01, **p<0.005 and ***p<0.001.

This latest result was also reflected in the spatial position of individual centroids (panel B Figures 7 and 8 for temporal and spatial task respectively). For the right hemisphere of both tasks, in the majority of the tested subjects, the clusters of voxels selective to the shorter durations had centroids located more anteriorly (see the y axis, diamonds in warmer color) with respect to the voxels responsive to the longer durations (diamonds in colder color). In the both hemispheres there was no significant difference in the spatial progression (wRD) of the vertices between the two tasks (temporal vs spatial task: left hemisphere p=0.427, right hemisphere p=0.520).

When we considered the preferred durations at the group level, we found for both tasks and both hemispheres that voxels lying closer to the posterior border of the chronomap preferred the longer durations, whereas those lying furthest preferred the shortest duration (Panel C Figures 7 and 8 for time and space task, respectively).

Within the IPS, we did not find a clear topography, neither at the group nor at the single subject level (see Supplementary Figure 10).

**Figure 10.**
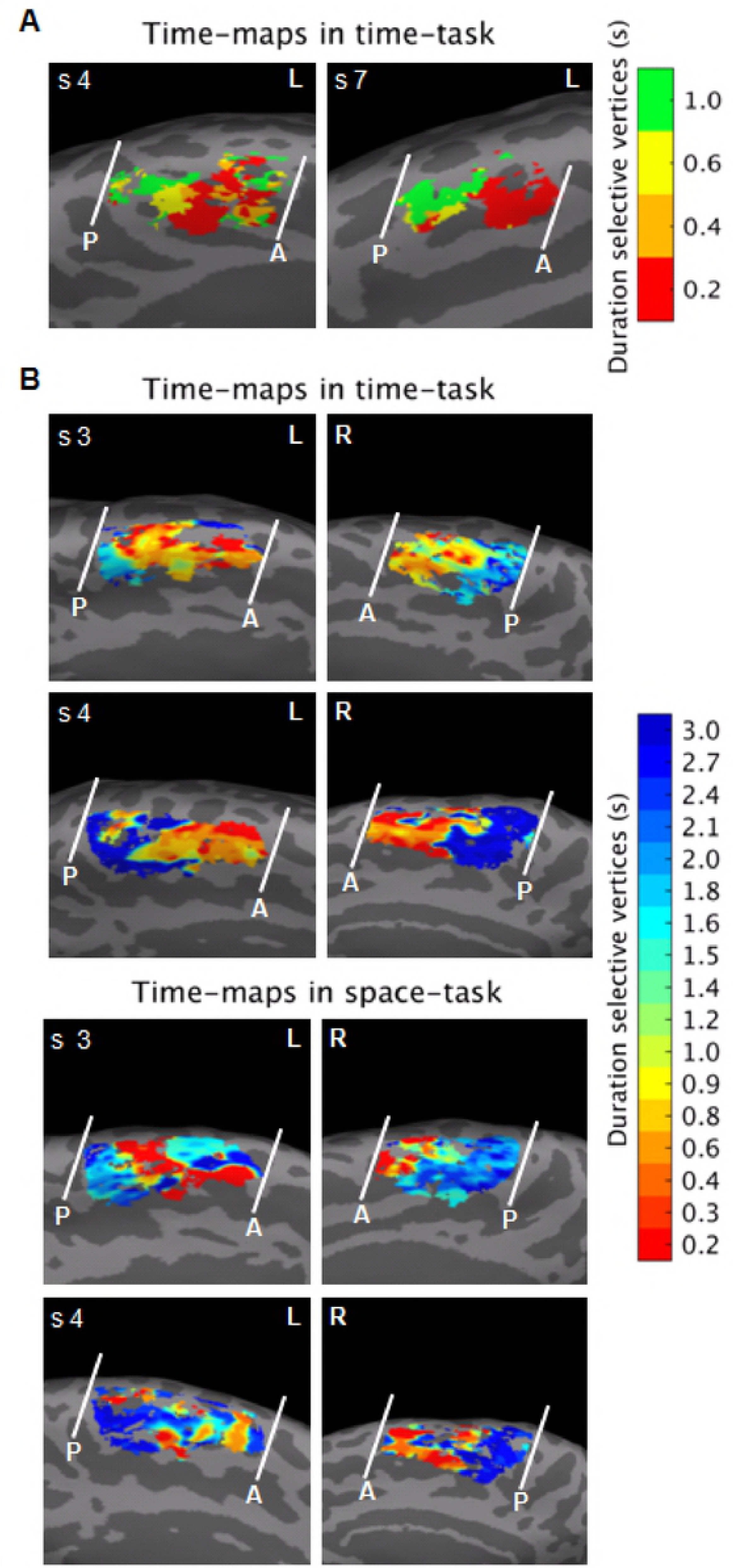
fMRI results, individual data of Exp.1 (A) and Exp.2 (B). (A) For 2 subjects of Exp.1 we show the left SMA chronomap with the anterior (A) and posterior (P) borders. Individual maps were obtained using a winner-take-all procedure based on statistical t-maps (T>3.13). We computed 4 different t-maps for each of the 4 S1 durations (p_FWE_-cluster level < 0.05, corrected for multiple comparisons across the whole brain). For the maps of whole sample (N=11) of subjects see Supplementary Figure 2. (B) For 2 subjects of Exp.2 we show the left and the right SMA chronomap with the anterior (A) and posterior (P) borders in the time (leftward) and in the space (rightward) task. The maps were the results of a pRF analysis. Here we show the projection on the cortical surface (medial part of BA 6) of the 17 estimated *µ* parameter. The 17 *µ* are the 17 durations presented (S1 when is either the standard or the comparison duration) in the 10 different trial types. Different colours represent vertices selective to different duration ranges (i.e., vertices with different estimated *µ*). For the maps of whole sample (N=10) of subjects see Supplementary Figures 6-9.

<u>Please Figure 7 and 8 here</u>

To examine the response tuning of duration sensitive voxels, also in this second experiment, we looked at the variation of the hemodynamic response as a function of the presented duration i.e., preferred versus non-preferred durations. Figure 9 shows the normalized hemodynamic response of SMA duration selective voxels to preferred and neighboring durations (PD and PD ± 1, see darker shades) as opposed to the response to distant durations (PD ± 2, see lighter shades). Given the limited number of repetitions for each of the 17 presented durations, for the plot of the signal change, we grouped the durations according to the 10 different trial types (i.e., 10 pairs of durations). The normalized BOLD response is plotted for both the time (upper panel) and the space task (lower panel). The bar plot shows that for the majority of duration selective voxels activity was enhanced for preferred and neighboring durations and suppressed for more distant durations (see Figure 9). Since there was no difference in the tuning analysis of left and right hemispheres, the plot shows the average tuning of left and right SMA.

<u>Please Figure 9 here</u>

Given the limited number of trials for each of the 17 presented durations, for this plot we grouped the durations selective voxels according to the 10 different trial type. On the x-axis are the 17 presented durations grouped in 10 different duration ranges. The BOLD signal in duration selective voxels is aligned to the presentation timings of the different duration ranges (i.e., 2^nd^ volume after S1 offset). The colored diamonds represent the point in time where the hemodynamic response of duration selective voxels matched the presentation timing of the appropriate duration (e.g., red-labeled voxels when the shortest range of duration is presented).

## Discussion

To summarize, here we showed with two independent data sets and two different paradigms and methods of data analysis, the existence of neuronal units tuned to different durations in SMA. Duration selectivity had a clear topographical organization in the rostro-caudal direction for, respectively, short and long durations. Chronotopic maps were observed across a wide range of durations (from 0.2 to 3 s) and not only at the group level, but also with a certain degree of variability at the single-subject level. Figure 10 shows for Exp.1 (panel A) and Exp.2 (panel B) the SMA chronomaps in two “ideal” subjects i.e., subjects with an anterior-short to-posterior-long spatial progression. This progression was present in 7 out of 11 subjects in Exp.1 (left SMA) and in 9 out 10 subjects in Exp.2 (right SMA, see Supplementary Figures 2, 7 and 9 for the SMA maps of all subjects).

Chronotopic maps were also task independent; maps were indeed found when time was available but it was task irrelevant. At tuning level, we found that the hemodynamic response in duration selective voxels was enhanced for preferred and neighboring durations and suppressed for durations far from the preferred one.

<u>Please Figure 10 here</u>

Neuronal tuning is an encoding mechanism widely used in neurons to represent sensory and motor information[13][19] and even more abstract features like quantities[14]. This topographic organization is thought to have a computational benefit, for example the efficiency of neural communication[20].

Duration selective cells have been previously reported in monkeys’ medial premotor cortex[3][4]. The present study extends this representational format to humans and shows that duration-selective units in this region are topographically organized along the anterior-to-posterior axis. Moreover, while the presence of duration-selective units in monkey’s premotor cortex was exclusively associated with motor timing behavior, our study shows the presence in human premotor cortex of duration-selective mechanisms in a purely temporal perceptual task.

In humans, duration selective mechanisms have been recently suggested by an fMRI study showing duration adaptation effects in the activity of the inferior parietal lobule (i.e., the Supramarginal Gyrus)[11]. Activity in this region is suppressed when consecutive stimuli have the same duration.

Our data support this finding and show the presence of duration selective mechanisms in a closer location i.e., the IPS, although in the left rather than the right hemisphere. However, our data go beyond this previous finding by showing a) the existence of duration selective activity for a wider range of durations, b) duration selectivity not only in the IPS but also in the SMA and c) most importantly we showed that only activity in the SMA is topographically organized in a way that neuronal units selective to similar durations occupy contiguous portion of the cortical surface so as to form chronomaps.

Moreover, similarly to the repetition suppression shown by Hayashi and colleagues in the SMG, the chronomaps in SMA were also present in the spatial task, when time was available but was task-irrelevant.

The presence of topography in SMA, but not in IPS, may indicate that duration selectivity in different brain regions (IPS and SMA) serves different purposes along the process leading to duration judgments.

Our hypothesis is that duration selective activations in premotor cortex may reflect an active reconstruction of temporal signals coming from different regions of the brain (e.g. visual or parietal areas)[21][2][22]. One can think of chronomaps in SMA as a temporal read-out, a later stage of duration encoding in which duration information becomes finally available and decision-making takes place. The IPS duration selectivity, which lacks a clear topography [11], may represent an intermediate stage where duration signals coming from low-level sensory regions are automatically organized. A support to this hypothesis comes from the observation that the perturbation of right SMG activity via Transcranial Magnetic Stimulation (TMS) affects time representations in the SMA[22].

Another element, in line with the idea that the duration tuning and topography observed here do not represent a low-level stage of temporal processing, i.e., something equivalent to sensory maps, is the anatomical location of the maps. Chronomaps were mainly observed in SMA and neither in the parietal nor in sensory regions. SMA has been implicated in a variety of timing tasks[16][23][24] with a range of durations spanning from a few hundreds of milliseconds to a few seconds[25][26] and with stimuli from different sensory modalities[27][28][29]. It is therefore likely that this area constitutes an ‘amodal’ and ‘high-level’ core of a timing network in which duration is represented in an abstract form independent of specific sensory modality or motor behavior.

Duration selective units were maximally responsive to the preferred duration, activated by neighboring durations, and exhibited the strongest suppression to durations distant from the preferred one. This seems to suggest a Gaussian-like type of response profile, where neuronal units tuned to similar durations have overlapping tuning curves. This tuning profile is also in line with the behavioral effects obtained with duration adaptation paradigms where an optimal proximity between “adaptor” and test duration leads to stronger repulsive effects[9]. In analogy with spatial vision or audition (e.g. visual orientation[13] or auditory pitch[30]), the tuning profiles observed here may serve the function of enhancing the discriminability of durations by suppressing the activity for different durations.

In summary, here we found a topographic representation of time in human premotor cortex, an area that has been previously identified as “time” region. Our findings of chronomaps clarify the nature of duration information represented there and, most importantly, indicate duration tuning and topography as possible mechanisms for duration read-out.

## Materials and Methods

### Subjects

We tested a total of twenty-one healthy volunteers, eleven in Exp.1 (5 females, mean age 23.7 years, SD 4.3 years) and ten in Exp.2 (9 females, mean age 27.7 years, SD 5.1 years) with normal or corrected-to-normal vision. All volunteers gave written informed consent to participate in this study, the procedures of which were approved by the ethics committee of the Faculty of Biology and Medicine at the University Hospital of Lausanne.

### Stimuli and Procedure

In Exp.1, we used a temporal discrimination task of visual durations. Visual stimuli were sinusoidal Gabor patches (100% contrast, spatial frequency of 1.9 cycles/degree, Gaussian envelope with standard deviation of 2.2 degrees, diameter of ∼9 degree) with a circular hole (diameter 0.6 degrees, at the center of the Gabor) displayed at the center of the screen around a central fixation spot (a black disk 0.5 degrees of diameter at a viewing distance of 90 cm) on a grey background. In each trial, two Gabor patches (S1 and S2) were sequentially presented with a variable inter-stimulus-interval ranging between 4 and 5.2 s in 0.08 s steps. The two stimuli were followed by a response cue i.e. a red fixation spot of 2 s duration (see Figure 1A). S1 and S2 varied in orientation and duration, although only duration was task relevant. The duration of S1 could be 0.2, 0.4, 0.6, and 1 s and its orientation 36, 72, 108, and 144 degrees. S2 could be either shorter or longer in duration than S1. The duration of S2 was longer or shorter by a constant Weber ratio of 0.4 (e.g. if S1 was 0.2 s, S2 was either 1.6 or 3.6 s), whereas the orientation of S2 was a value randomly chosen from the 4 possible orientations used for S1 (i.e., 36, 72, 108, or 144 degrees). The combination of duration and orientation lead to 16 different types of S1 stimuli. Each stimulus type for S1 was presented only once in each fMRI run.

Participants were asked to judge whether the duration of S2 was shorter or longer than S1. Participants made their responses by pressing one of two buttons on a response-pad. They used their right index finger to express the choice “S2 shorter than S1” and their right middle finger for the “S2 longer than S1” responses. Participants were instructed to be as accurate as possible (no emphasis was put on reaction times) and to fixate at the center of the screen while performing the duration discrimination task. They were also requested to ignore the orientation changes of the stimulus and to not use counting strategies to estimate duration.

Each fMRI run contained 16 trials and the total duration of each run was 3 min and 51 s. We collected 18 fMRI runs in two separate sessions (9 runs per session). The second session was performed 1–3 days after the first session. The data of this first experiment are partially shared with another study that is currently under review (Hayashi et al.).

In Exp. 2, two tasks were used: a temporal discrimination and an orientation discrimination task. The stimuli and the task structure were identical in the two tasks; the only difference was the stimulus feature participants were asked to attend (duration versus orientation). The stimulus was a sine wave grating (size = 400 by 400 pixels, 8.01 degree of visual angle at viewing distance of 90 cm; spatial frequency was 0.05 cycle/pixel), drifting at a speed of 1 cycle per second and displayed at varying angular orientations. Within a trial the sequence of orientation changes was fixed and was always leftward first, rightward second and vertical last (Figure 1C). Within the two ‘main orientations’ (leftward - rightward) the grating continuously changed its orientation at a rate of 5 Hz (an orientation change each 0.2 s) and the range of changes was between 30° and 45°. The amount of time the grating maintained its ‘main’ orientation defined a temporal interval. During the temporal discrimination task, participants judged which of the two ‘main orientations’ (leftward or rightward) was maintained for a longer time. In the orientation discrimination task participants judged which of the two ‘main orientations’ underwent the biggest change. In this manner, the physical stimuli were identical and the amount of attention paid to them was equated across tasks, the only difference was the instruction given to the participants (attend to duration versus attend to orientation changes). The vertical orientation signaled the time to make the response (by pressing one of two response keys on a keypad) and it was also the inter-trial-interval. The duration of the vertical orientation was kept constant (1.37 s), whereas the duration of the two ‘main orientations’ varied.

On each trial there was always a standard (T) and a comparison duration (T+ΔT). The duration of the comparison was a constant proportion of the standard (i.e., 50% of the standard, Weber ratio was equal to 0.50). The presentation order of standard and comparison (i.e., standard first, comparison second or vice-versa) was randomized and counterbalanced across trials. Half of the times S1 was a standard and the other half it was a comparison duration. We used 10 different standard durations, ranging from 0.2 to 2 s in steps of 0.2 s, one for each trial. The full combination of standards and comparisons resulted in following 10 pairs of durations 1: 0.2-0.3 s, 2: 0.4-0.6 s, 3: 0.6-0.9 s, 4: 0.8-1.2 s, 5: 1.0-1.5 s, 6: 1.2-1.8 s, 7: 1.4-2.1 s, 8: 1.6-2.4 s, 9: 1.8-2.7 s, 10: 2.0-3.0 s.

While the grating was displayed for a standard and a comparison duration, its angular orientation changed at a rate of 5 Hz. The angular change was one of 12 pseudo-randomly chosen values ranging from 30° to 45° (in logarithmic steps, base 10). It is worth emphasizing here that since the orientation changes were chosen pseudo-randomly, sometimes the same orientation could be displayed more than once (maximum number of allowed repetitions of the same orientation was 3). Therefore, the number of orientation changes was not entirely predictive of the duration of the stimulus.

The differences between rightward and leftward orientation could be 5°, 7°, 9° or 11°. We chose these different values based on the results of a purely behavioral pilot study where we tested both temporal and orientation discrimination tasks. The angular differences chosen were those leading to discrimination accuracy similar to the temporal task.

Both tasks were structured in ‘ascending’ and ‘descending’ cycles. Each cycle comprised 10 trials and lasted 44 s. ‘Ascending’ cycles started with the shortest duration pair (i.e., 0.2-0.3 s, first trial) and ended with the longest pair (i.e. 2-3 s, the tenth trial). On descending cycles, it was the reverse (i.e. the first trial had the longest and the tenth the shortest pair). The time interval between cycles was 2.03 s; during this interval the grating was in vertical orientation. In both tasks subjects were responding using either the index or the middle finger of their right hand. In each fMRI run there were 10 cycles. There were separate runs for ‘descending’ and ‘ascending’ cycles (1 run each) and for the temporal and the orientation discrimination tasks (2 runs each). Each participant thus performed a total of 4 fMRI runs (220 fMRI volumes each).

### Behavioral Data Analysis

In Exp.1 for each participant we took the percentage of performance accuracy for the four different S1 durations and we entered these values in a one-way repeated measures ANOVA.

In Exp.2 for each participant we took the percentage of performance accuracy for the 10 different duration pairs in the two tasks and submitted them to a task (time, space) × durations (10 durations pairs) within subject ANOVA.

For both experiments the alpha level was set to 0.05. As post-hoc test we used the Bonferroni test.

### MRI Acquisition and Analyses

MRI Acquisition

The mapping of the selectivity of the neural responses necessitated high spatial resolution of the functional data. The increased signal-to-noise ratio and available BOLD associated with ultra-high magnetic field systems (>3 T) allowed the use of smaller voxel sizes in fMRI[31]. In addition, the spatial specificity of the BOLD signal is improved because the signal strength of venous blood is reduced due to a shortened relaxation time, restricting activation signals to cortical gray matter[31]. Therefore, we employed high-resolution, 7T fMRI for the functional maps.

In both experiments blood oxygenation level-dependent (BOLD) functional imaging was performed using an actively shielded, head-only 7T MRI scanner (Siemens, Germany), equipped with a head gradient-insert (AC84, 80 mT/m max gradient strength; 350 mT/m/s slew rate) and 32-channel receive coil with a tight transmit sleeve (Nova Medical, Massachusetts, USA).

In Exp.1 time-course series of 169 volumes were acquired for each run using the 3D-EPI-CAIPI sequence[32]. The spatial resolution was 2.0 mm isotropic, the volume acquisition time was 1368 ms, the flip angle was 14 degrees, the repetition time (TR) 57 ms and the echo time (TE) 26 ms and the bandwidth 2774 Hz/Px. The matrix size was 106 x 88 x 72, resulting in a field of view of 210 (AP) x 175 (RL) x 144 (FH) mm. An undersampling factor 3 and CAIPIRINHA shift 1 were used. Slices were oriented transversally with the phase-encoding direction left-right. 42x45 reference lines were acquired for the GRAPPA reconstruction. For each individual, a total of 3,042 volumes (169 volumes per run, 18 runs) were analyzed.

High-resolution whole-brain MR images were also obtained using the MP2RAGE pulse sequence optimized for 7T[33] (voxel size = 1.0 x 1.0 x 1.0 mm, matrix size 256 x 256 x 176, TI_1_/TI_2_ =750/2350ms, α_1_/α_2_ = 4/5 degrees, TR_MP2RAGE_/TR/TE = 5500/6.5/2.84 ms).

In Exp. 2 fMRI data were acquired with a continuous EPI pulse sequence with sinusoidal read-out (1.5 × 1.5 mm in-plane resolution, slice thickness = 1.5 mm, TR = 2000 ms, TE = 25 ms, flip angle = 47°, slice gap = 1.57 mm, matrix size = 148 × 148, field of view 222 × 222 mm, 40 oblique slices covering most of occipital, parietal and premotor regions). In each fMRI run we acquired 220 fMRI volumes. A T1-weighted high-resolution 3D anatomical image (resolution = 1 × 1 × 1 mm, TR = 5500 ms, TE = 2.84 ms, slice gap = 1 mm, matrix size = 256 × 240, field of view = 256 × 240) was acquired for each subject using the MP2RAGE pulse sequence. For each participant an additional whole-brain EPI image (a single volume with 80 slices and TR=4000ms and otherwise identical parameters to the functional data) was acquired in order to aid the co-registration between the EPI images and the individual MP2RAGE. The EPI sequence used in Exp.2 did not allow whole-brain coverage. Based on the results of Exp.1, we chose to place the 6-cm thick imaging slab so as to cover the occipital, parietal and premotor cortices.

fMRI Analyses

fMRI Preprocessing

For both experiments, functional imaging data were preprocessed using the statistical parametric mapping toolbox (SPM12, Wellcome Department of Imaging Neuroscience, University College London). In Exp. 1 the EPI volumes acquired in each session were realigned to the mean of the session and then co-registered to the T1-weighted image acquired in the same session. In order to perform group level analysis (see Figure 2) the realigned and co-registered images were then normalized to the averaged DARTEL template (diffeomorphic anatomical registration through exponentiated lie algebra[34]) and smoothed with a 2 mm full-width at half-maximum Gaussian kernel. To perform surface-based analysis, data were kept in the subject’s space i.e., after realignment and co-registration to the T1-weighted image data were then directly smoothed with a 2 mm full-width at half-maximum Gaussian kernel (Figures 3A and 4).

In Exp. 2 the EPI volumes acquired in each session were slice time corrected, realigned to the mean of the session and then co-registered first to the whole brain EPI image and subsequently to the T1-weighted image acquired in the same session. Since the sequence used for Exp.1 was a 3D-EPI-CAIPI (i.e., the whole k-space was acquired at once, with no time lags), only in Exp.2 data were slice time corrected. In order to performed volumetric analyses and to visualize the group-level pRF results a DARTEL temple was also created for Exp.2.

### General Linear Model (GLM) Analysis

Exp.1 data were analyzed using a GLM approach. The fMRI time series were first analyzed in each single subject. Each single subject model included 18 runs/session with 6 event-types in each session. These comprised the 4 different S1 durations (each event was time-locked to the offset of S1), a fifth event time locked to the onset of S2 (comparison duration) and a sixth event time-locked to the onset of the participants’ response. The linear models included also the motion correction parameters as effects of no interest. The data were high-pass filtered (cutoff frequency = 0.0083 Hz). In order to see brain activity correlated to the different S1, for each subject we estimated 4 contrasts, one for each S1. These contrasts also averaged parameter estimates across the 18 runs.

In order to test the existence of chronomaps in the group, the four contrast images estimated in each subject, were then entered into a second-level ANOVA where we performed again 4 different contrasts (one for each S1 duration). The statistical threshold was set to p < 0.05 FWE cluster-level corrected for multiple comparisons across the entire brain volume (cluster size estimated at a voxel level threshold p-uncorrected = 0.001).

Correction for non-sphericity[35] was used to account for possible differences in error variance across conditions and any non-independent error terms for the repeated-measures.

To appreciate the existence of chronomaps, the 4 t-maps, obtained either at single subject or at group level were then used to classify the voxels according to their preference to one of the 4 different duration ranges. Voxels were classified according to a “winner take all” rule, for example voxels with the greatest t value (threshold was set to T> 3.13) for the shortest duration range (0.2 s) were classified as responsive to that duration range and labeled with number 1. We created 4 different labels and each label was associated with a specific color for visualization purposes.

### pRF Analysis

Data from Exp.2 were analyzed using the Population Receptive Field method (pRF). The pRF analysis was performed with the SamSrf toolbox for pRF mapping (https://figshare.com/articles/SamSrf_toolbox_for_pRF_mapping/1344765/22).

This toolbox implements a method of analysis similar to the one used in several studies[14][17][18][36]. We performed the pRF analysis on two distinct ROIs: BA6 and IPS. The ROIs were based on the Freesurfer software’s Broadmann and Destrieux atlases. (http://surfer.nmr.mgh.harvard.edu/). For each subject the pRF analysis was performed on slice-time corrected, realigned, co-registered and smoothed images.

The idea behind pRF is that neuronal receptive fields are a form of tuning functions that reflect specific stimulus properties. For each subject pRFs were modeled as one-dimensional Gaussians with 2 parameters: *µ*, the stimulus duration and the spread of the pRF, σ. For the pRF modelling we used the offset of all S1 durations, no matter whether S1 was a standard or a comparison duration. This procedure led to the identification of 17 durations (i.e., 0.2. 0.3, 0.4, 0.6, 0.8, 0.9, 1, 1.2, 1.4, 1.5, 1.6, 1.8, 2, 2.1, 2.4, 2.7, 3.0 s). For each time point (i.e. each TR) of our fMRI timeseries and each vertex of the ROIs, the method estimates the overlap between the Gaussian tuning model of a given *µ* and the presented durations. A coarse-to-fine optimization approach then determined the optimal pRF parameters for which the goodness-of-fit of the predicted time series to the observed data was maximized. The maps shown are the projection on the cortical surface of the estimated optimum *µ* parameter. Different colors represent vertices (i.e., voxels projected onto the cortical surface) selective to different duration ranges.

For the group-level analysis, the pRF maps for each participant were morphed into a common DARTEL template using the morph labels feature of the MNE software (https://mne-tools.github.io/dev/index.html). MNE performs the morphing between subjects using the spherical surfaces provided by Freesurfer.

### Visualization

For visualization of the group and of the single subject fMRI results in both experiments we inflated either the DARTEL template (group level results) or the single subject T1-weighted image (individual results) using the FreeSurfer pipeline (http://surfer.nmr.mgh.harvard.edu/). To reconstruct surfaces for the DARTEL template, the gray matter and white matter images of the template were combined into a single image with two distinct values assigned to the gray matter and white matter voxels. The combined images were treated as a skullstripped T1-weighted image and submitted to the Freesurfer pipeline for surface reconstruction.

### Quantification of the spatial distribution of chronomaps

Surface-based approach

In order to better appreciate the spatial distribution of the chronomaps at the individual level, we identified chronomaps in each single subject by using either the single-subject SPM t-maps (Exp.1) or the pRF maps (Exp.2). The surface-based analyses were performed on images in the subject’s space.

For a better visualization, these volumetric maps were projected onto the cortical surface of each individual brain. Individual cortical surfaces were reconstructed following the Freesurfer pipeline via segmentation of different brain tissues (projection fraction was set to 0.5).

Individual chronomaps were identified in left SMA and left IPS for Exp.1 and left and right SMA, and left and right IPS for Exp.2. In Exp.1, we used anatomical landmarks (i.e., identification of the pre, post-central gyri and intra-parietal sulcus) to make sure that chronomaps at single subject level matched the location of those observed at group level.

For Exp.2 the identification of the chronomap at single subject level was easier since the pRF analysis was performed on 2 distinct ROIs: BA6 and IPS. For the identification of SMA chronomap we took only the medial part of the BA6.

For each map we created a surface-ROI (left SMA and left IPS for Exp.1 and left, right SMA and left, right IPS for Exp.2) and we manually draw its borders. According to the spatial progression of the maps (from short to long duration-selective voxels) observed at group level, we identified an anterior and a posterior border for SMA maps and a medial and a lateral border for IPS. Those borders were then used as cuts to flatten the surfaces and became the outer edges of the flattened surfaces. For SMA we took the postcentral gyrus as anatomical landmark for drawing the posterior border.

For each duration selective vertex and each ROI, we calculated the weighted Relative Distance (wRD) from one of the borders of the map (D_1_). This border, arbitrarily chosen, was the posterior for SMA map, and the lateral for the IPS map.

In more detail, the weighted Relative Distance from D_1_ border was computed as following: 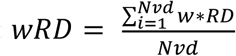, where is the weight of each vertex defined as the ratio between clustered duration-selective vertices (*Nbrs*) and the total number of vertices maximally responsive to a given duration (*Nvd*) i.e., *w* = *Nnbrs*/*Nvd*. Whereas *RD* was the ratio between the distance from one of the borders (*D*) and the mean distance between the two borders (*TD*) *RD* = *D*_1_/*TD*.

For each map we computed the wRD of each duration selective vertex and we identified a slope of the spatial progression of those vertices. The individual slopes were used to perform a Wilcoxon test in order to check the statistical significance of the spatial progression of the maps.

### Volume-based approach

To make sure that the results from the surface-based analyses depicted reality and were not the product of wrong projection of voxels onto the surface, we also performed volume-based analyses. The analyses on the volume were performed on data normalized to the Dartel space i.e., Dartel-11 (Exp. 1) and Dartel-10 (Exp. 2). Similarly to surface based analysis, also here we identified for each experiment and each subject chronomaps in SMA and IPS.

Also, for volumetric maps we defined maps’ borders. These were anterior and posterior for SMA, and medial and lateral for IPS.

In order to check whether the duration selective voxels followed the same spatial progression as the surface-maps, we identified for each subject and each map the “preferred duration” of different portions of the map. More precisely, we binned the individual volumetric ROIs in parallel planes of 1.5 mm width. Within each volumetric-bin the “preferred duration” was calculated as the duration the majority of activated voxels responded to. Thus, for each subject, we had a sequence of preferred durations between the two borders of the map. We then decided to compute the average of preferred durations across subjects. Since different subjects had sequences of preferred durations of different length, we decided to proceed as follow: we calculated for each spatial bin its relative distance from one border (D1) of the map (i.e., posterior for SMA and lateral for IPS). Then for each map we created a single sequence of “preferred durations”, which included the sequences of all subjects ordered according to their relative distance from D1. In order to reduce the total length of this long sequence, we averaged every five values of the sequence. The result of this procedure is displayed in panel C of Figures 3, 7 and 8.

In order to appreciate the spatial distribution of the maps at single subject level, for each subject and each duration-selective cluster of voxels we also estimated the “weighted Centroids” (wCntrs). Within a cluster of duration selective voxels, every voxel was assigned a weight based on the number of neighboring voxels with the same duration selectivity. This means that clustered voxels had more weight than sparse ones. The wCntrs were then calculated by taking into account the position of all duration-selective voxels within a cluster but each position was represented as many times as the weight assigned to a specific voxel. This measure allowed us to visualize in a single graph the central position of all duration selective clusters in all subjects (see panel B, Figures 3, 7 and 8).

### Tuning Analysis

To check the response properties of duration selective voxels we looked at the BOLD response to preferred and non-preferred durations. In both experiments, for each subject and each cluster of duration selective voxels within the different chronomaps (i.e., SMA and IPS in the left hemisphere for Exp.1 and left and right SMA for Exp.2), we looked at the normalized hemodynamic response to preferred and non-preferred durations.

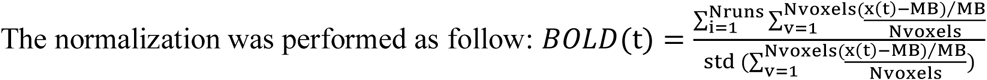

where t is the signal in a given voxel and MB is the baseline obtained averaging the signal of t across runs. Normalization was then performed subtracting the signal in a given voxel from a baseline value and dividing it by the baseline. The BOLD response was aligned to the 2^nd^ volume (i.e., a TR) after the offset of the S1 duration. Within a single subject, we first averaged the BOLD signal across the voxels of a cluster and then across the fMRI runs.

## Acknowledgments

Financial support has been provided by the European Research Council -ERC (Grant Agreement No 682117 BiT-ERC-2015-CoG to D.B.), the Swiss National Science Foundation (Grants #3100B0-133136 and 320030_149982 to M.M.M.), the Japan Science and Technology Agency (JST to R.K) and JSPS Postdoctoral Fellowships for Research Abroad and Research Fellowships for Young Scientist (to M.J.H). We would like to thank Dr. Mayur Narsude for supporting data collection.

## Author Contributions

RK and DB conceived the study; WvdZ, DB, MJH collected the data; FP, SK, RK, GB, and DB analyzed the data; RK, FP, MJH, MMM, WvdZ, and DB wrote the paper.

